# Biochar alters soil properties, microbial community diversity, and enzyme activities, while decreasing conifer performance

**DOI:** 10.1101/2021.05.17.444392

**Authors:** Jake Nash, Jessica Miesel, Gregory Bonito, Monique Sakalidis, Han Ren, Daniel Warnock, Lisa Tiemann

## Abstract

Biochars are porous charcoal-like materials that can enhance soil health and plant growth, but its use has not been adequately evaluated in woody cropping systems. We set up an experimental Christmas tree plantation on a Marlette series soil amended with two types of biochar and conducted two studies on the impacts of biochar on the agroecosystem over three years following establishment. The first study investigated the effects of biochar on plant performance, soil physicochemical properties and extracellular enzyme activities, while the second investigated the response of the root-associated fungal community. Both biochars stimulated five extracellular enzyme activities, with increases of between 67% and 446%. Structural equation modelling identified increases to dissolved organic carbon and soil moisture as potential mechanisms of biochar’s effects on enzyme activities. Tree growth and survival were negatively affected by biochar application, depending on the tree species and biochar applied, which may have been due to induced nitrogen limitation. High-throughput sequencing showed that biochar decreased the diversity of root-associated fungal communities, with the ectomycorrhizal species *Wilcoxina mikolae* reaching levels of hyper-dominance on balsam fir in response to one of the biochars. Further studies should investigate how biochar can be harnessed to remediate specific soil quality issues or restructure soil ecosystems in ways that improve crop performance.

## 1. Introduction

Biochar is a porous charcoal soil amendment produced by pyrolyzing plant or animal biomass (Lehmann and Joseph, 2009). Because of its relative chemical recalcitrance, biochar acts as a stable carbon (C) sink and has been proposed as means to offset anthropogenic CO_2_ emissions (Lehmann et al., 2006). Biochars have highly variable physical and chemical properties depending on the feedstock material and manufacturing specifications (Enders et al., 2012; Gundale and DeLuca, 2006; Sohi et al., 2010). The totality of a specific biochar’s physical and chemical properties determines its effect (or lack thereof) on soil properties including pH, water-holding capacity, cation exchange capacity, bulk density, and nutrient availability (Dai et al., 2020; Gundale and DeLuca, 2007; Jeffery et al., 2011). Biochar also has been shown to impact plant- and soil-associated microbial communities (Lehmann et al., 2011; Zhang et al., 2018). In an ideal scenario biochar can significantly increase plant resistance to pathogens and climate change induced stresses by increasing the benefits of associating with root colonizing fungal symbionts and growth-promoting bacteria (Ali et al., 2017; Ishii and Kadoya, 1994; Matsubara et al., 2002; Robertson et al., 2012).

Under certain circumstances, biochar’s effects on soil physicochemical properties and plant-microbe interactions can result in increased plant growth and crop yields (Major et al., 2010; Robertson et al., 2012; Rondon et al., 2007). Although past work has demonstrated the strong potential for biochar to improve crop performance, the overwhelming majority of studies have been in annual cropping systems and much less is known about how biochar may affect crop performance and soil biology in woody, perennial cropping systems. There have been a number of studies demonstrating neutral or negative effect of biochar on plant performance and plant-soil interactions (Biederman and Harpole, 2013; Bonanomi et al., 2017; Elzobair et al., 2016a; Gaskin et al., 2010; Hagner et al., 2016; Lehmann et al., 2011; Van Zwieten et al., 2010; Warnock et al., 2007; Warnock et al., 2010). These studies highlight the context-dependency of the effects of biochar application on plant-soil relations and demonstrate the necessity of understanding the outcome of specific biochar-crop-soil combinations as a prerequisite to widespread usage. Many of the benefits of biochar are likely to be the most pronounced in sandy, well-drained soils, which are the most prone to drought and nutrient leaching. In well-drained sandy soils, biochar could be effective at reducing irrigation costs during increasingly frequent periodic droughts by increasing soil water-holding capacity (Ali et al., 2017). Furthermore, biochar could decrease requirements for fertilizer application by limiting nutrient losses from leaching out of sandy soils and stimulating the ectomycorrhizal (ECM) fungi that associate with Christmas trees, which can increase the nutrient-acquisition capabilities of their plant hosts (Lehmann et al., 2011; Robertson et al., 2012; Warnock et al., 2007).

Soil microorganisms mineralize organic matter by secreting extracellular enzymes including phosphatases, aminopeptidases, cellulases and chitinases, which liberate smaller and more labile forms of C and nutrients (Burns et al., 2013; Grandy et al., 2008; Sinsabaugh et al., 2008). Once liberated by the action of extracellular enzymes, the more labile forms of C and nitrogen (N) are readily available for uptake by plants and soil biota. Lab incubation and greenhouse studies have already shown positive effects of biochar on soil extracellular enzyme activities (Al Marzooqi and Yousef, 2017; Elzobair et al., 2016a; Masto et al., 2013; Paz-Ferreiro et al., 2014; Paz-Ferreiro et al., 2012). These studies indicate that biochar amendments may represent a useful approach for increasing soil microbial community activity rates and thus for increasing soil health and crop yields (Luo et al., 2018). However, as with the plant growth and soil fungal responses mentioned above, instances of neutral and negative effects of biochars on individual enzyme activities have also been reported (Elzobair et al., 2016b; Lammirato et al., 2011; Paz-Ferreiro et al., 2012). Despite the frequently observed increases of extracellular enzyme activities (EEAs) in response to biochar, few studies have attempted to elucidate the mechanisms of biochar’s effects on EEAs (Bailey et al., 2011; Khadem and Raiesi, 2017; Khadem and Raiesi, 2019). A mechanistic understanding of biochar’s effects on EEAs could help predict which types of biochar will result in desirable changes to EEAs.

We established an experimental Christmas tree plantation featuring two commonly growth Christmas trees, blue spruce (*Picea pungens*) and balsam fir (*Abies balsamea*) and conducted two separate, but related, studies investigating the effect of two different conifer wood-derived biochars on soil, plants, and microbes. The first study tested the hypotheses that the biochars would 1) stimulate extracellular enzyme activities, 2) ameliorate unfavorable soil properties including nutrient concentrations and moisture, and 3) improve plant performance. The second study tested the hypothesis that biochar would modify root-associated fungal communities and increase ectomycorrhizal diversity and abundance.

## 2. Methods

### Experimental setup

#### 2.1 Site description

We established the field experiment in May of 2016 in a previously fallow field at Michigan State University’s Tree Research Center, in Lansing, Michigan, USA (42°40’28.9”N 84°28’01.8”W). Soils are Marlette series (fine-loamy, mixed, semiactive, mesic Oxyaquic Glossudalfs) and slopes range from 0% to 3.0%. Mean annual air temperature at the field site is approximately 18°C, the mean annual rainfall is 80.5 cm and the average annual snowfall is 130 cm. The herbaceous layer is dominated by non-native weeds, including *Setaria viridis* (green foxtail), *Solidago* spp. (goldenrod) and *Conyza canadensis* spp. (horseweed).

#### 2.2 Biochar manufacturing processes and characteristics

Our study evaluated one biochar from each of two different Michigan based companies, Biogenic Reagents (BGR) and US Biocarbon (USB). The USB biochar was made from waste wood pallets (southern yellow pine species) and produced in a continuous carbonizer at 550°C for 18 min. BGR biochar was produced from woody pine biomass residue (woodchips, sawdust, limbs, etc.) from forestry operations in the upper peninsula of Michigan (primarily *Pinus resinosa* and *P. banksiana*) via pyrolysis at 650°C for 30 minutes in a rotary reactor system. Carbon (C) and nitrogen (N) contents of the biochars were determined on an elemental analyzer (Costech Analytical Technologies, Inc., Valencia, CA, USA) while the concentrations of other elements in the surface layer of the biochars were determined by scanning electron microscopy with energy dispersive X-ray spectroscopy taken on three replicate subsamples (described in detail in section 2.5; see table 1 for biochar properties).

#### 2.3 Study design and establishment

The experimental design included biochar application two rates as well as a control (0 Mg ha^−1^, 25 Mg ha^−1^, and 75 Mg ha^−1^; hereafter referred to as the control, low rate, and high rate, respectively) and two species of popular Christmas trees – balsam fir and Colorado blue spruce. To implement these treatments in a fully factorial design, we divided the study site into two units, each of which was further subdivided into six columns of 30m length and 1m width, with 1m wide buffer strips between columns for a total of 12 treatment areas (see Fig. S1 for a map of study layout). Three columns in each unit were planted with each species. Each treatment area consisted of fifteen 2m x 1m plots, each of which was planted with three tree seedlings. We applied the biochars on May 8 (BGR) and May 9 (USB), 2016 using a turf spreader followed by discing to a depth of 6” to incorporate the biochar. We then transplanted three bareroot seedlings into each plot with 0.5 m between seedlings within each plot. This planting rate yielded an initial count of 540 seedlings. The close spacing provided extra seedlings that could be destructively harvested while leaving the remaining seedlings at a typical spacing (2 m) for Christmas tree plantations (Landgren et al., 2003). To control competition from weeds, we applied the herbicides Simazine, Flumioxazin, Glyphosate, Clethodim, Pendimethalin, and 2,4-D at various points throughout 2016, 2017, and 2018, which are described in detail in table S1. Although herbicides can affect microbial processes, they were applied evenly over the study area and were thus unlikely to have introduced bias into our results We irrigated seedlings during periods of drought with a tractor sprayer. We did not fertilize the experiment or apply any soil amendments other than the biochar. We used the same field experiment to conduct two separate but related studies on the effects of biochar on the agroecosystem – study 1 investigated the effects of biochar on soil properties, microbial nutrient cycling, and plant performance, while study 2 focused on the assembly of root-associated fungal communities. The trees that we destructively sampled to examine root-associated fungal communities in study 2 were taken from separate plots than those used in study 1 to avoid disturbing soil processes. Because of this, we did not attempt to relate the soil data from study 1 with the fungal community data from study 2.

### Study 1: Effects of biochar on soil properties, enzyme activities, and plant growth

#### 2.4 Field sampling

We sampled soil multiple times per year during growing seasons (June through October) in 2016, 2017, and 2018 by taking a minimum of five independent samples from a subset of plots within each treatment combination, yielding 50 total samples for most sampling timepoints (5 biochar treatments x 2 tree species x 5 replicates). Dates of soil sampling as well as the analyses that were performed on each set of samples are listed in table S2. Each of the five independent samples from within each treatment consisted of three cores (0 to 15 cm depth) from beneath the canopy of a single seedling within the sampled plots that were combined into one soil sample to reduce variability stemming from fine-scale spatial heterogeneity in soil properties. We also installed cation and anion plant root simulator (PRS) probes (Western Ag, Saskatoon, SK, Canada) in five plots per treatment from July 17, 2018 to September 13, 2018 to measure the bioavailability of the following 15 ions over the growing season – NO_3_, NH_4_, Ca, Mg, K, PO_4_, Fe, Mn, Cu, Zn, B, S, Pb, Al, and Cd. PRS probes bind ions in soil solution and provide an integrated index of the availability of those ions throughout the deployment period.

We quantified weed biomass by clipping all herbaceous plant (weed) tissues at the soil surface within a 0.25 m^2^ quadrat. We separated plant shoots into grasses and forbs, dried them to a constant weight and then weighed samples for the total dry mass of grasses and forbs. We conducted surveys of the growth and survival of seedlings in the Spring and Fall of 2016, 2017, and 2018. Seedling surveys recorded seedling mortality, root collar diameter, and height. We calculated a biomass proxy for each seedling by multiplying the basal area (cm^2^) by the height (cm). This metric approximates the volume of the central stem of each seedling.

#### 2.5 Laboratory soil analyses

Soil properties that we assumed to be dynamic (dissolved nutrients, microbial biomass, enzyme activities) were generally measured at multiple timepoints and soil properties that we expected to be relatively stable (bulk density, total C/N) were measured at a single timepoint, as outlined in table S2. Following sampling, we sieved soils to 5 mm to allow larger biochar particles to pass through and stored them at 4° C for a maximum of one week before freezing them at −20°C or processing them for analysis. We sent soil samples collected shortly after planting in 2016 to the soil testing service at the University of Maine (Orono, Maine, USA) to determine soil P, K, Mg, S, pH, and effective cation exchange capacity. We determined total soil C and N concentration via dry combustion (Costech ECS 4010 CN analyzer; Costech, Valencia, CA USA). We measured soil bulk density by taking 5.0 cm x 15.0 cm soil cores with an impact corer, weighing the volume of soil sampled, and using a subsample to measure soil moisture. For soil bulk density, we sampled five cores across each biochar treatment, combining the plots planted with spruce and fir trees because we did not expect tree species identity to affect soil structure after only three seasons. We determined soil microbial biomass, dissolved organic carbon (DOC), total dissolved nitrogen (TDN), NO_3_, and NH_4_ on soil samples taken from multiple dates in 2016, 2017, and 2018 (see table S2). We prepared soil extracts by shaking 8 g of field-fresh soils with 40 mL 0.5m K2SO_4_ at >250 rpm for 60 minutes and then filtering them through Whatman #1 filter paper. We stored filtrates at −20° C until analysis. We used the chloroform fumigation-extraction method (Vance et al., 1987) to determine soil microbial biomass C and N, with both fumigated and unfumigated extracts analyzed for DOC/TDN on a varioTOC analyzer (Elementar, Langenselbold, German). Microbial biomass C and N were calculated as the differences in extractable C and N in fumigated and unfumigated extracts, using efficiency factors of 0.45 and 0.54, respectively (Brookes et al., 1985; Jenkinson et al., 2004). We quantified soil NO_3_ concentration using a modified version of the enzyme reduction method described in detail by Wittbrodt et al. (2015), with absorbance measured at 540 nm on a Biotek synergy H1 microplate reader (Winooski, VT, USA). We assessed soil NH_4_ contents by using the microplate protocol described in Sinsabaugh et al. (2000).

We used microplate-format colorimetric and fluorimetric assays (Saiya-Cork et al., 2002) to determine seven different soil EEAs involved in C, N, and P cycling – β-glucosidase (BG), cellobiohydrolase (CBH), leucine aminopeptidase (LAP), β-N-acetyl glucosaminidase (NAG), peroxidase (PER), phenoloxidase (PHEN), and acid phosphatase (PHOS), with fluorescence and absorbance measured on a Biotek synergy H1 microplate reader (Winooski, VT, USA). Briefly, we prepared soil slurries by homogenizing 1 g (wet weight) of soil in 125 mL of water with an immersion blender. We chose to suspend soil samples in water rather than a buffer to allow assays to proceed at the soil’s natural pH (German et al., 2011). We used methylumbelliferyl-linked substrates for BG, CBH, NAG, and PHOS assays, a methylcoumarin-linked substrate for the LAP assay, and L-DOPA as a substrate for PHEN and PER assays. All fluorimetric assays were run with substrates at 40 μM concentration. We incubated assays for approximately 18 hours. We included homogenate blanks, substrate blanks, and buffer blanks in all assays.

We investigated whether biochar underwent physical or chemical changes following application by using scanning electron microscopy with energy dispersive X-ray spectroscopy (EDX). We recovered biochar particles from soil cores sampled in October 2018 by dispersing ~15 mL of soil in 400 mL of deionized water and picking them out of the water with forceps. This method was intended to remove loosely adhering debris, while allowing tightly adhering soil particles to remain. We also selected biochar particles that had not been applied to the field and had been stored for three years to represent samples without field aging. We prepared biochar particles for mounting by drying them at 60° C for two days, after which we mounted them to stubs with quick curing epoxy and coated them with osmium (≈10 nm thickness) in an NEOC-AT osmium chemical vapor deposition coater (Meiwafosis Co., Ltd., Osaka, Japan). We performed scanning electron microscopy using either a JEOL 6610LV or JEOL 7500F scanning electron microscope (JEOL Ltd., Tokyo, Japan) equipped with an Oxford Instruments AZtec system (Oxford Instruments, High Wycomb, Bucks, England) for EDX. We performed EDX on unaged biochars and on portions of aged biochars that were free of encrusted soil particles to avoid confounding the elemental composition of the soil with that of the biochar. We used Michigan State University’s Center for Advanced Microscopy for all sample preparation and imaging.

#### 2.6 Statistical analyses

For variables that were measured at multiple timepoints, we tested the effects of biochar application using repeated measures mixed models with the nlme package in R (Pinheiro et al., 2013). We inverse or log transformed data to meet the assumptions of normality when absolute values of skewness exceeded two or those of kurtosis exceeded seven (Kim, 2013). We used pairwise t-tests or Mann-Whitney U tests (when data could not be transformed to meet assumptions of normality) to evaluate the significance of differences between biochar treatments and the control at individual sampling dates and did post-hoc tests that averaged the effects of biochar application across all sampling dates using the emmeans package in R (Lenth et al., 2018).We used chi-square tests to test for differences in tree survival between treatments. Because pairwise tests were only performed between the control and each of the biochar treatments, we calculated Bonferroni corrections by multiplying P-values by four (2 biochars x 2 rates = 4 comparisons). After finding strong effects of biochar on EEAs and identifying soil physicochemical properties that were affected by biochar, we used structural equation modelling to test which soil physicochemical characteristics could be identified as potential mechanisms of biochar’s effects on EEAs. In the structural equation models, we accounted for the effect of sampling date by first fitting linear models to EEAs and all soil physicochemical variables with sampling date as the only explanatory variable. We used the residuals from these models in all structural equation models so that we could test for relationships between variables while controlling for variation due to sampling date. We conducted all structural equation modelling with the package lavaan in R (Rosseel, 2012). We identified modifications to DOC, inorganic N, and soil moisture *a priori* as potential mechanisms of biochar’s effects on EEAs because these variables were correlated with both biochar application and EEAs for at least one of the biochars. We used these three physicochemical variables in structural equation models with paths leading from biochar to these variables, and also from these variables to EEAs. We also included a direct path from biochar to EEAs. We calculated net coefficients representing the importance of each mechanism of biochar’s effects on EEAs by multiplying the coefficient leading from biochar application to each soil physicochemical property and the coefficient leading from each soil physicochemical property to each enzyme. We calculated these coefficients as standardized coefficients which vary from −1 to 1. We used bootstrapping to calculate the standard errors and to test for significance of these coefficients. In total, we fit ten different structural equation models to the data to test for the effects of the two biochars on the five hydrolytic enzymes. We did not include oxidative enzymes in the structural equation models because they were not significantly affected by biochar application.

### Study 2: Effects of biochar on root-associated fungal communities

#### 2.7 Sequencing of root-associated fungal communities

We profiled root-associated fungal communities by performing amplicon sequencing of the fungal ITS region. In late October of 2017, we destructively sampled seedlings from across all treatments (N = 5) and froze intact root systems at −80°C until processing. We prepared root samples for DNA extractions by thawing them, washing them free of loosely adhering soil, and then drying them at 35°C. Tightly adhering soil was not washed from the root system and was included in the extraction because it is likely to contain rhizosphere and root-associated fungi. Dried root samples were gently crushed by hand in a paper bag and fine roots that easily broke off from the root system were collected and ground by hand between two sheets of paper into a fine powder (Benucci et al., 2016). We extracted DNA from 50 mg of ground root material using the DNeasy PowerSoil Pro Kit (Qiagen, Hilden, Germany) following the manufacturer’s instruction. We prepared libraries for sequencing using a three step PCR that has been described previously (Benucci et al., 2019; Chen et al., 2018) with the primers ITS1F and ITS4 (Gardes and Bruns, 1993; White et al. 1990). In the first step, we used the primers without adapters to amplify the ITS region for ten cycles. In the second step, we used primers modified with frameshifts and Illumina adaptors for ten cycles of PCR (Lundberg et al., 2013). In the final step, we ligated barcodes and sequencing adaptors with 15 PCR cycles. We normalized PCR products to a concentration of 1-2 ng/μl using the SequalPrep Normalization Plate Kit (Thermo Fisher Scientific, United States) and were then pooled in equal volumes. We used Agencourt AMPure XP magnetic beads to remove small fragments and primer dimers (Beckman Coulter, United States). We sequenced samples on an Illumina MiSeq at Michigan State University’s Genomics Core Facility.

#### 2.8 Sequence Analysis and Statistics

We used FastQC to quality control sequences (Andrews, 2010) and demultiplexed sequences in QIIME 2 (Bolyen et al., 2019). We removed primers and conserved regions from sequences using Cutadapt v2.6 and USEARCH v10 (Edgar, 2016; Edgar and Flyvbjerg, 2015; Martin, 2011). We clustered sequences into operation taxonomic units (OTUs) at a 97% similarity threshold using the UPARSE algorithm of USEARCH (Edgar, 2013) and discarded singletons. We assigned taxonomy to representative sequences using the naïve Bayesian “feature-classifier” plugin in QIIME 2 (Bokulich et al., 2018; Bolyen et al., 2019) trained on the February 2, 2020 release of the UNITE database (Kõljalg et al., 2005). We assigned taxonomically identifiable OTUs to guilds using FUNGuild (Nguyen et al., 2016). We conducted downstream analyses on the OTU matrix using the phyloseq and vegan packages in R (Ihaka and Gentleman, 1996; McMurdie and Holmes, 2013; Nguyen et al., 2016). We performed alpha and beta diversity analyses on two different OTU matrices. In the first OTU matrix, representing the full dataset, we rarefied samples to an even sequencing depth of 13472, which excluded three samples with lower sequencing depth from analysis. In the second OTU matrix, we filtered to only include OTUs which were designated as ectomycorrhizal (ECM) by FUNGuild. We then calculated the relative abundance of ECM OTUs as their proportion among all ECM OTUs without rarefaction. We used this second dataset to test for effects on the composition and diversity of the ECM community specifically. We calculated the Shannon and Simpson diversity indices and the number of observed OTUs as measures of alpha diversity, which we tested for significance with ANOVA and pairwise t-tests with Bonferroni corrections. We calculated beta diversity with the Bray-Curtis dissimilarity metric, plotted with principal coordinates analysis (PCoA), tested for homogeneity of variance, and tested for significance with permutational multiple analysis of variance (PERMANOVA). We tested for effects of tree species and biochar treatment on the abundance of individual taxa that occurred in at least 40% of samples in each comparison using analysis of composition of microbiomes (ANCOM) on unrarefied, non-relativized OTU matrices, with a W statistic cutoff of 0.6 used for significance (Mandal et al., 2015). For all analyses on the fungal community data, we combined the low and high application rates for each biochar into a single grouping to increase statistical power. Representative sequences, OTU tables, OTU taxonomy assignments, and sample metadata are available in supplemental files 2-6.

## 3. Results

### Study 1: Effects of biochar on soil properties, enzyme activities, and plant growth

#### 3.1 Soil physicochemical properties

The application of BGR biochar at the highest rate resulted in strong and persistent increases in dissolved organic C (DOC, Fig. 1a; ANOVA, *P* < 0.0001), with significant effects observed at four out of six sampling dates (t-test, *P* < 0.0001), and an increase of 58% averaged across all sampling dates (pairwise test, *P* <0.0001). At the lower application rate, BGR biochar increased DOC by 9% averaged across all sampling points (pairwise test, *P* = 0.07), while USB biochar treatments did not alter DOC. Biochar had less clear-cut effects on total dissolved nitrogen (TDN), which demonstrated either increases, decreases, or a lack of an effect in response to biochar treatments, depending on the sampling date (Fig. 1b). Across all sampling dates, there were no consistent effects on biochar on TDN. Together, these two patterns show that BGR biochar increased the DOC pool without modifying the TDN pool, which led to predictable increases in the DOC:TDN ratio at both application rates (pairwise test, low rate *P* = 0.02, high rate *P* < 0.0001). Microbial biomass C and N were little affected by biochar treatment. At the lower application rate, BGR biochar increased microbial biomass N relative to the control (pairwise test, *P* = 0.03), but this effect was not observed at the higher application rate of the BGR biochar or in the USB biochar treatments.

**Figure 1.**
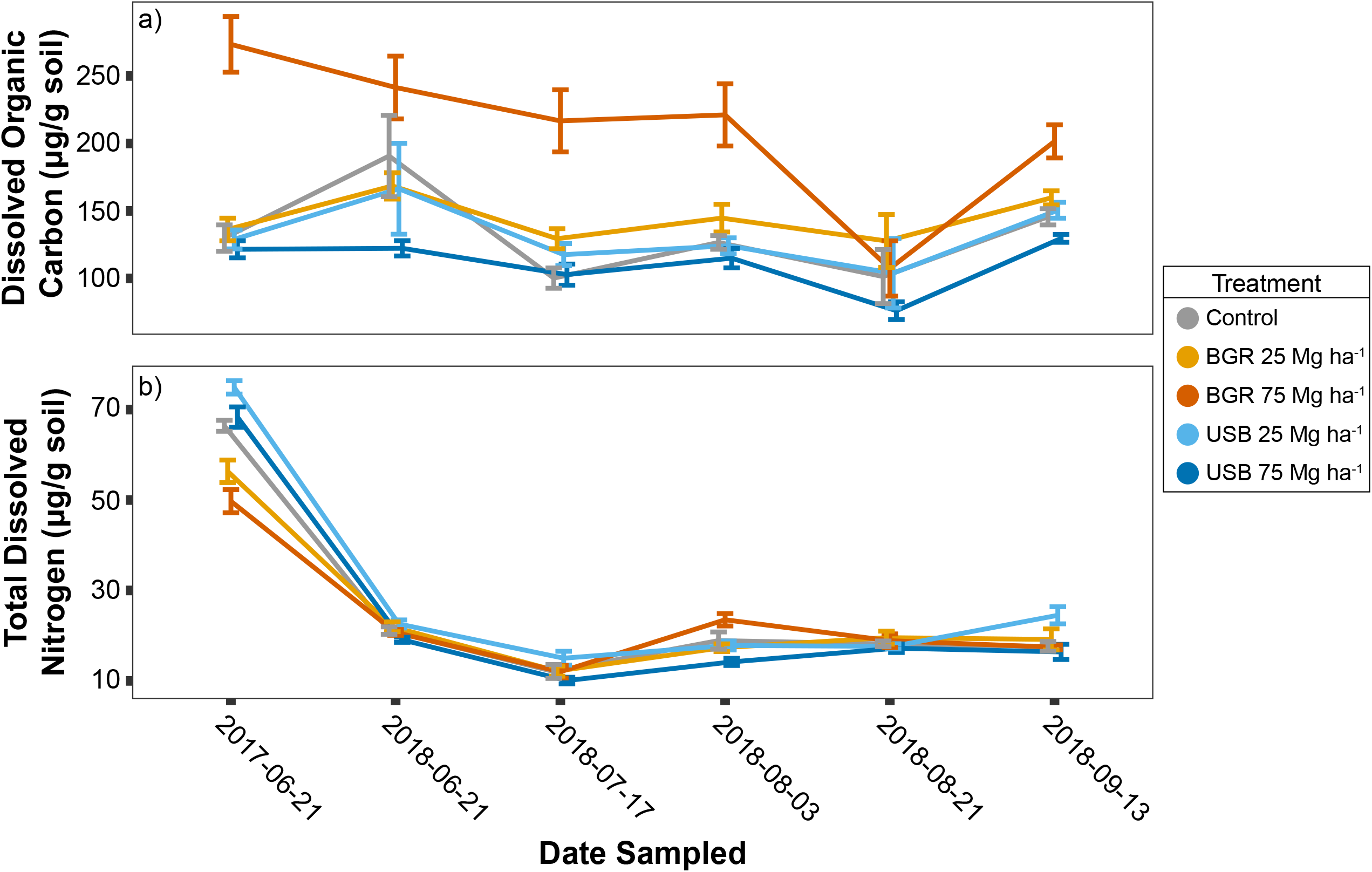
Mean concentrations and standard errors of dissolved organic carbon (a) and total dissolved nitrogen (b) are plotted in each treatment group at six sampling dates in 2017 and 2018 with the standard error displayed. Note that the location of points along the x-axis are offset slightly to minimize overlap and improve legibility.

Generally, both types of biochar positively affected soil moisture content in 2016 (ANOVA, *P* <0.0001), 2017 (*P* < 0.0001), and 2018 (*P* < 0.0001), with greater effects at the higher application rate (see Fig. S2). Although USB-amended plots generally had higher soil moisture than control plots, pairwise t-tests for each sampling date were never significant while the effects of BGR on soil moisture were much more pronounced and often significant, especially at the higher application rate (see Fig. S2). However, these effects were not entirely consistent across sampling dates. For example, in 2016 there was a significant interactive effect between biochar treatment and sampling date on soil moisture (ANOVA, *P* = 0.017), with negative effects of USB low rate (*P* = 0.034), USB high rate (*P* = 0.036), and BGR low rate (*P* = 0.006) on July 29. This sampling date came five days after a rainstorm in which 22 mm of precipitation fell in a single day. These negative effects of biochar on soil moisture content contrast with positive effects of both types of biochar on soil moisture that were observed on the day after the same storm (July 25). Throughout 2017 and 2018, there were not significant negative effects of any biochar treatments on soil moisture, though on some dates there was an absence of any positive effect.

In addition to its effects on soil moisture, biochar modified a number of soil physicochemical properties. At the end of the first growing season, both biochars had increased soil Mg and P, while BGR biochar at the higher application rate had increased Ca, K, B, Mn, Na, S, and effective cation exchange capacity (pairwise t-test, Bonferroni *P* < 0.05). USB biochar decreased Al and Fe contents (pairwise t-test, Bonferroni *P* < 0.05). Total soil N was not affected by biochar application two months after the establishment of the experiment (July 19, 2016), though we did not analyze soils for total N at later timepoints and cannot rule out longer term effects. BGR biochar decreased soil bulk density by 10.1% (*P* = 0.099) and 20.7% (*P* < 0.001) relative to the control treatment at the low rate and high rate respectively, while USB biochar had no effect on bulk density at either application rate (Bonferroni-corrected *P* = 1). Two months post-application, biochar had resulted in shifts in soil pH from 5.81 (in the control plots) up to 6.72 (*P* = 0.007) and 7.12 (*P* < 0.0001) at the high rate for USB and BGR biochars respectively with weaker, though still significant effects at the low rate (see Fig. S3).

Plant root simulator (PRS) resin strips installed in 2018 showed that both types of biochar decreased the bioavailability of NO_3_, Fe, Mn, Cu, and Pb and increased the bioavailability of NH_4_ and K (ANOVA, *P* < 0.05) during the 2018 growing season (Fig. S4). Although biochar increased the bioavailability of NH_4_, the vast majority of the resin-bound inorganic N was in the form of NO_3_, which exhibited a strong negative response to biochar application, dwarfing the positive response of NH_4_. For almost all ions that were significantly affected by biochar application, effects of BGR biochar were greater than those of USB biochar (see Fig. S4). Effects of biochar on other ions were less consistent between biochar types. Resin-bound phosphate decreased at the USB low rate and increased at the BGR high rate, while Mg increased with USB and decreased with BGR at both application levels. The bioavailability of Ca, Zn, B, SO_4_, Al, and Cd were not affected by either biochar. Biochar decreased the bioavailability of the three heavy metals Fe, Cu, and Pb by around 56% and 84% for USB and BGR biochars, respectively, applied at the higher rate, although there was no effect on other heavy metals (Mn, Zn, and Cd). We collected data on extractable soil inorganic N data across 2016, 2017, and 2018 which largely support the patterns revealed by the PRS probes but also demonstrates the potential for sporadic disruptions to the general trend of lower N availability in the biochar treatments (see Fig. S5). Immediately following the establishment of the experiment (July 7, 2016) inorganic N content decreased significantly (*P* < 0.0001) in all biochar treatments (11.8-13.6 μg N/g dry soil) relative to the control (32.4 μg N/g dry soil). Treatment effects quickly became less pronounced, but still with generally lower concentrations of inorganic N (see Fig. S5). There was an exception to this pattern on August 21, 2018 when BGR-amended plots had greater soil inorganic N than control plots at both biochar application rates (*P* < 0.0001).

#### 3.2 Enzyme activities

Averaged across all sampling dates in the first growing season (2016), biochar induced significant increases to the soil extracellular enzyme activity (EEA) of BG, CBH, LAP, NAG and PHOS which were mostly observed in the high rate treatments of both USB and BGR biochar treatments. Yet this pattern was not consistent and there were significant treatment by date interactions for all enzymes (ANOVA), indicating temporal instability (see Fig. 2a-e). However, in the two following seasons (2017 and 2018), patterns in hydrolytic EEAs became much more stable, with positive and application rate-dependent effects of both biochars observed at almost all sampling dates (see Fig. 2f-k; ANOVA, *P* <0.0001). Across 2017 and 2018, effect sizes were generally greatest in the BGR high rate treatment, which resulted in increases of 95%, 67%, 446%, 83%, and 183% for BG, CBH, LAP, NAG, and PHOS activities respectively. Effects of the two biochars at the low rate on enzyme activities were not always significant, but effects at the high rate were highly significant for all hydrolytic enzymes in 2017 and 2018 (see Fig. 2f-k). This pattern was generally stable throughout these two years, with date by treatment interaction terms only significant for CBH (ANOVA, *P* < 0.0001) and PHOS (*P* = 0.02), although this was driven by changes in the relationship of activities between some of the BGR and USB biochar treatments, rather than by their relationship to the control treatment which almost always had the lowest mean EEA for the hydrolytic enzymes (see Fig. 2f-k). Among the five hydrolytic enzymes, effect sizes of biochar application were highest for LAP, averaging a 197% increase in activity in 2017 and 2018 relative to the control across the four biochar treatments. LAP was also the only enzyme that showed somewhat consistent increases in activity in response to biochar in 2016. There was never an effect of biochar on peroxidase activity, while phenoloxidase activity was negligible (often negative) and was not tested for significance.

**Figure 2.**
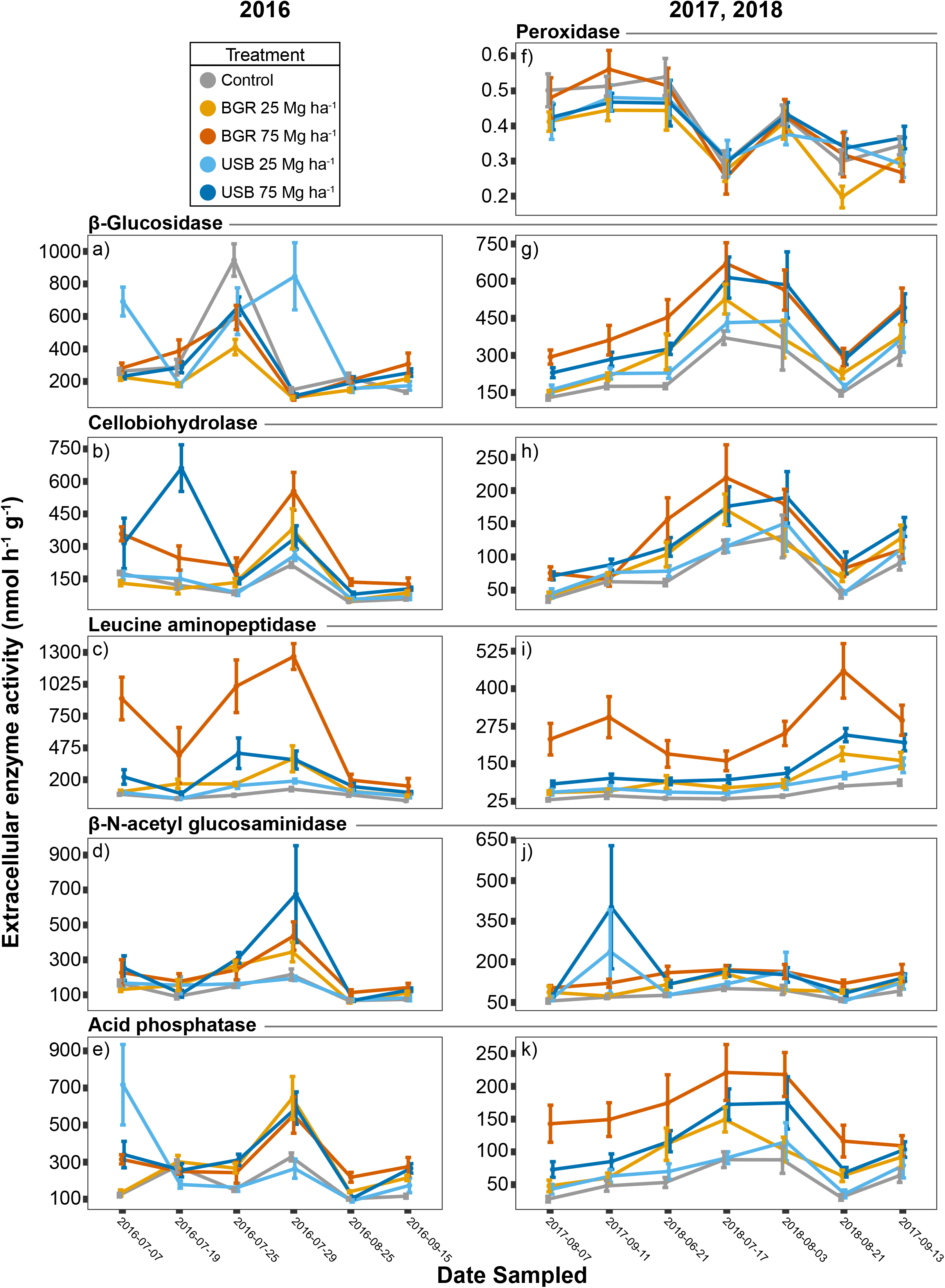
The mean extracellular enzyme activities (EEAs) of six enzymes in each of the treatments are plotted across 13 sampling dates in 2016 (a-e) and in 2017/2018 (f-k). Note that EEA data from 2016 and from 2017/2018 are plotted on different scales and that we do not have data on peroxidase activity from 2016.

Based on correlations that we observed in ANOVA models, we identified DOC, soil moisture, and inorganic N as potential mechanisms of biochar’s effects on EEA and further investigated their roles using structural equation modelling (see Fig. 3a, b for model structure). Structural equation modelling revealed that BGR biochar increased the activities of all five hydrolytic enzymes both indirectly by increasing soil moisture (which then stimulated EEA) and directly via unmeasured mechanisms (see Fig. 3a, c). BGR biochar also had indirect effects on leucine aminopeptidase and acid phosphatase (but not BG, CBH, or NAG) by increasing DOC content (see Fig. 3a, c). USB biochar had direct effects on all EEAs, but few indirect effects (see Fig. 3b, d) largely because the USB biochar had weaker effects on the soil variables that would have mediated such effects. There was however, a significant, weakly positive indirect effect of USB biochar on BG activity mediated by a negative effect on inorganic N (see Fig. 3d).

**Figure 3.**
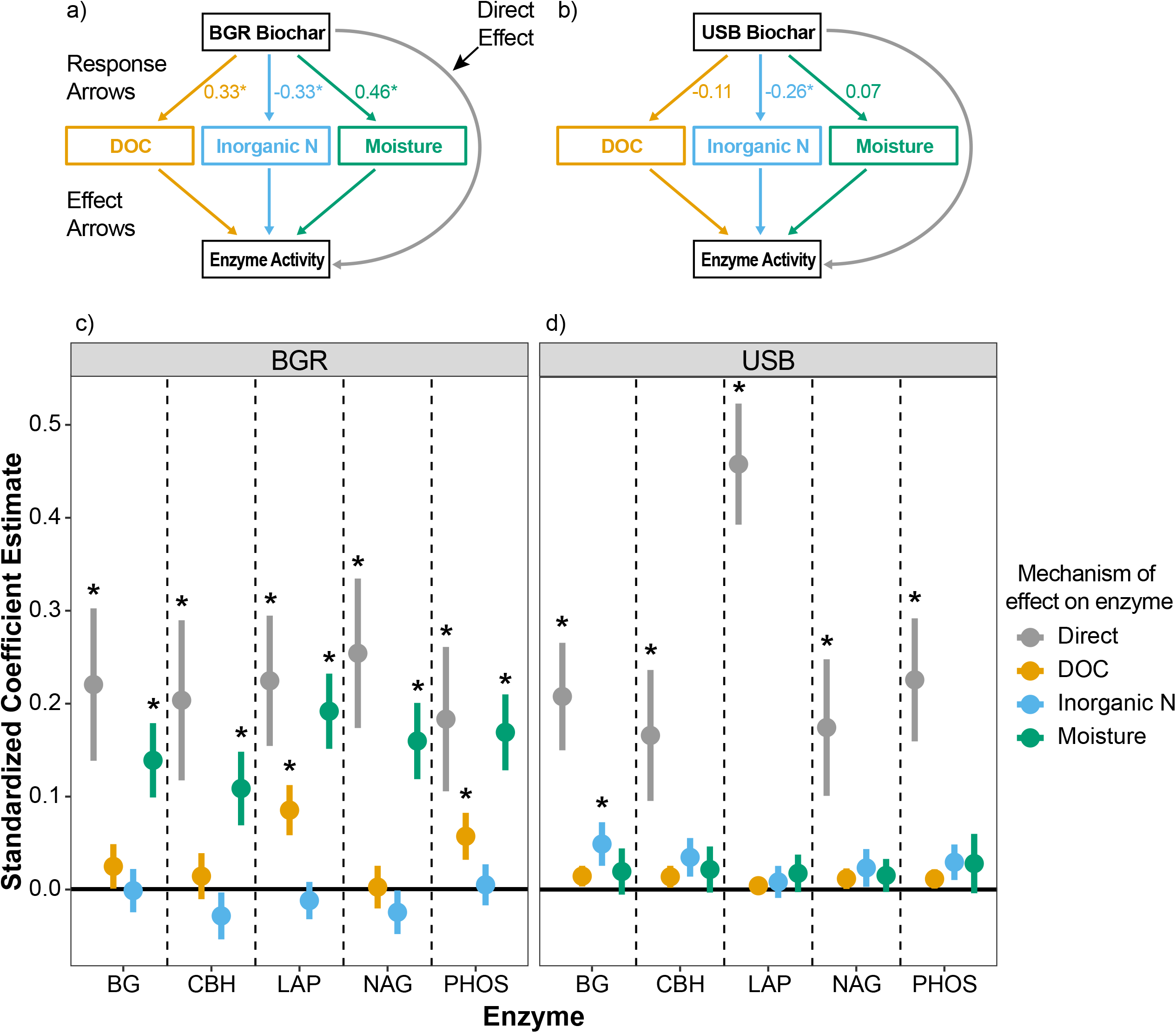
The results of ten structural equation models (5 enzymes x 2 biochars) that predicted extracellular enzyme activities (EEAs) based on direct effects of biochar and indirect effects mediated throughout effects on dissolved organic carbon (DOC), inorganic nitrogen, and soil moisture. (a,b) Model coefficients can be grouped into three categories (labelled in panel a) – response arrows (response of soil variables to biochar), effect arrows (effect of soil variables on extracellular enzyme activities), and the direct effect of biochar on extracellular enzyme activities. Standardized coefficients for the response arrows are displayed for each biochar, with asterisks indicating statistical significance at *P* < 0.05. Coefficients of the effect arrows are not displayed because these were different for each of the five enzymes. Mechanisms of biochar’s effect on extracellular enzyme activities are color-coded to match the plots in panels c and d. (c,d) Standardized coefficient estimates (with a range of between −1 and 1) of the direct and indirect effects of biochar on each of the five extracellular enzyme activities are plotted with standard errors displayed. The indirect effects were calculated by multiplying the coefficients of the response arrows by those of the effect arrows (marked in panel a). Asterisks indicate statistical significance of effects at *P* < 0.05.

#### 3.3 Electron microscopy investigation of biochar structure

In scanning electron microscope images, the microporous structure of the pine wood feedstock was readily apparent with visible pores and pits (see Fig. 4). Images taken on field-aged biochar showed that soil particles had encrusted areas of both types of biochar, often filling the pores (see Fig. 4 c, d), and that a filamentous fungus had colonized the surface of BGR biochar (see Fig. 4c), though it is unclear if this represents selective microhabitat usage or incidental colonization. There was no obvious evidence of physical degradation of biochar structure from visual assessment, and no substantial changes in the surface chemistry of either biochar after field aging (data not shown).

**Figure 4.**
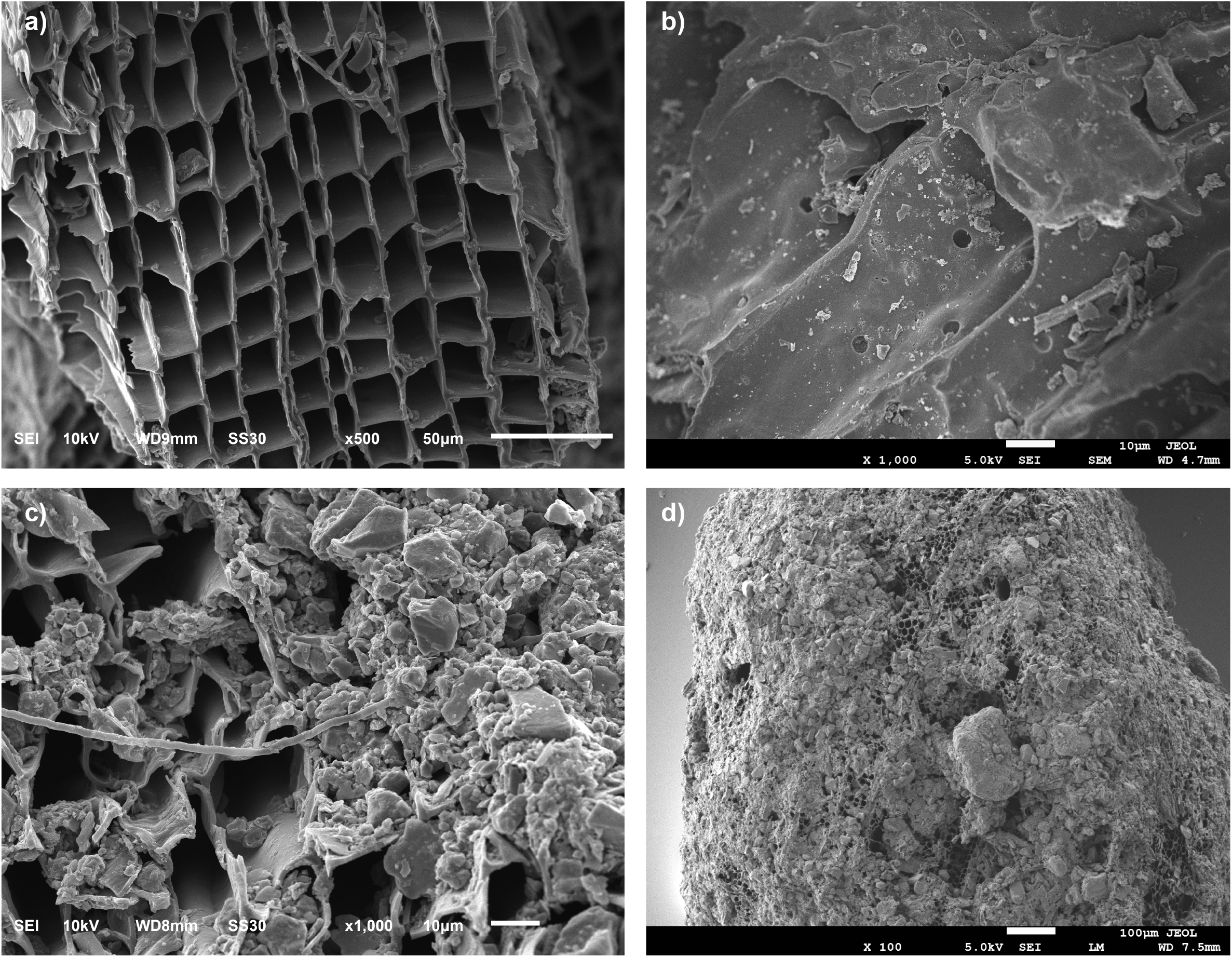
Scanning electron microscopy provides a glimpse into the interactions that biochar may have with soil and microbes. (a) a piece of BGR biochar that was recovered from field soil three years after application still had a relatively intact pore structure. (b) a piece of USB biochar still had noticeable pits in its xylem cell walls. (c) a piece of BGR biochar after three years in the field was crusted in soil particles and colonized by what appears to be a fungal hypha. (d) a piece of USB biochar after three years in the field was almost entirely covered in soil particles and may be serving as a nucleus for soil aggregation.

#### 3.4 Plant performance

BGR biochar induced significant mortality of spruce (see Fig. 5a) and fir (see Fig. 5b) trees and decreased the growth of fir (see Fig. 5d). For blue spruce, the BGR biochar low rate resulted in decreased survival that was immediately apparent by the first sampling date (chi-square test, *P* < 0.05) and persisted throughout the experiment, with survival of 7% compared to 38% in the control by the end of the experiment (Fall 2018). BGR biochar applied at 75 Mg ha^−1^ did not have an effect on blue spruce survival (see Fig. 5a). Balsam fir displayed decreased survival in both of the BGR treatments, starting in September of the first growing season (2016) and persisting throughout the experiment (chi-square test, P < 0.05, see Fig. 5b), with survival averaging 39% in the BGR treatments compared to 70% in the control treatment by the end of 2018. Balsam fir displayed lower growth in the BGR treatments beginning in the spring of 2017 (though it was sometimes only significant in the low rate treatment), with biomass proxy values in the BGR treatments approximately half of those in the control treatment by the end of the experiment (pairwise t-test, *P* < 0.05). There was generally no effect of USB biochar on the growth or survival of either tree species. In 2016, weed biomass was higher in all biochar treatments than in the control, though the only significant difference was with the BGR low rate which had 94% greater weed biomass than the control (pairwise t-test, Bonferroni *P* < 0.05). There were no significant differences in weed biomass in 2018.

**Figure 5.**
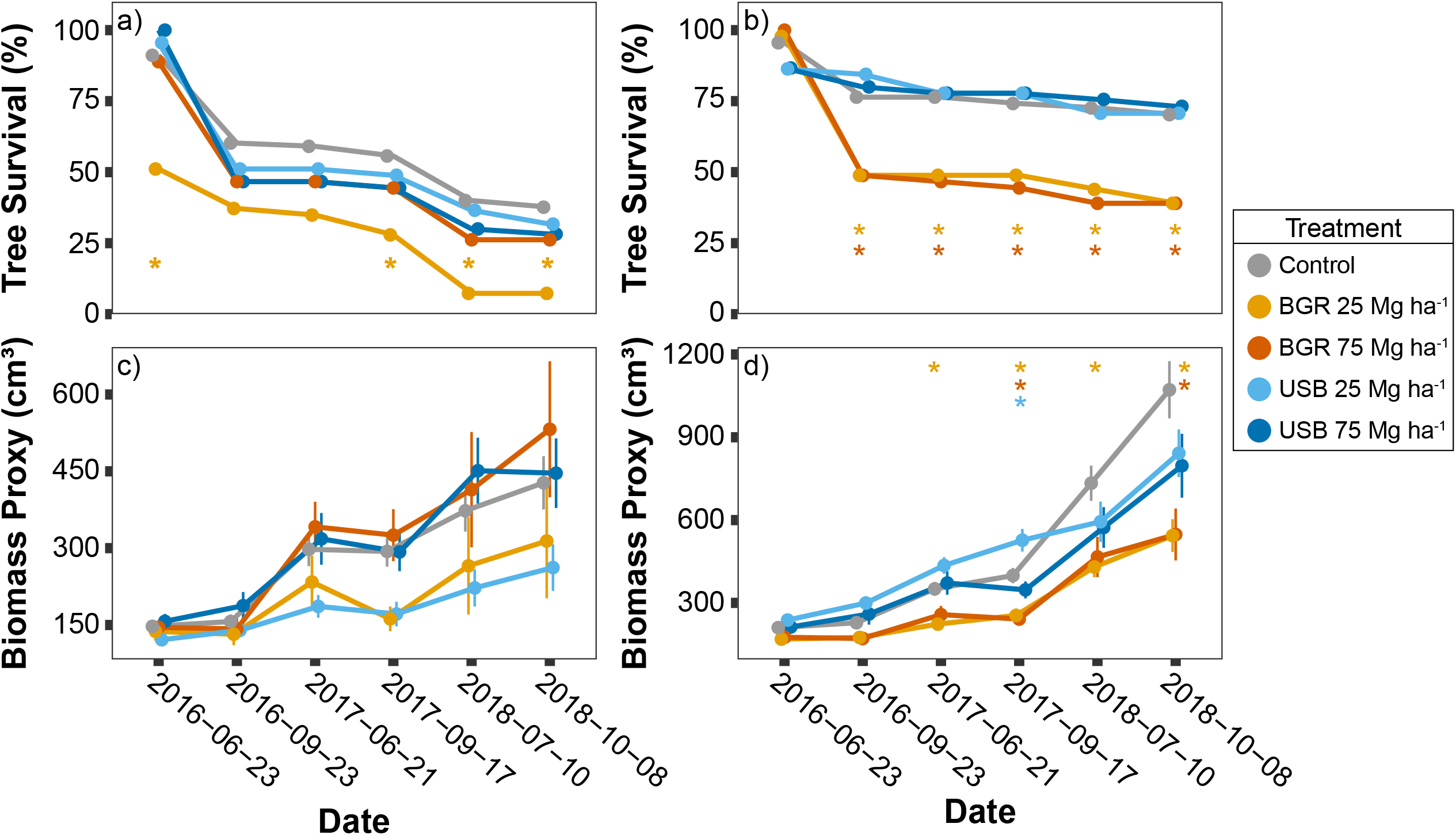
The percent surviving trees at six sampling dates across 2016, 2017, and 2018 is plotted for spruce (a) and fir (b). A proxy of tree biomass (calculated by multiplying tree basal area by tree height) is plotted for spruce (c) and fir (d) trees. In all plots, significant differences (Bonferroni-corrected *P* < 0.05) of biochar treatments from the control treatments is indicated by asterisks that are color coded to match the legend. Chi-square tests were used to test significant differences in tree survival and t-tests were used to test significant differences in the biomass proxy.

### Study 2: Effects of biochar on root-associated fungal communities

#### 3.5 Effects of biochar on fungal communities

Root-associated fungal communities had an average of 357 unique fungal OTUs per sample and 3,407 total OTUs across all 50 samples, although ectomycorrhizal (ECM) taxa constituted a small fraction of this diversity, with an average of only 6.4 ECM OTUs per sample and 51 total ECM OTUs (see Fig. 6a, b). Although ECM OTUs made up only 1.5% of the observed OTUS of the root associated fungal community, they constituted 7.5% of the total number of sequenced reads. ECM sequence relative abundance modestly increased in the USB biochar treatment relative to the control, though the effect was not significant (t-test, Bonferroni *P* = 0.14). For balsam fir-associated fungal communities, ECM OTU richness on balsam fir roots decreased from a mean of 10.6 OTUs in the control down to 4.8 and 5.4 OTUs in the USB and BGR biochar treatments respectively (ANOVA, *P* = 0.002; t-test, *P* = 0.01, 0.018), while total OTU richness decreased from a mean of 430 OTUs in the control down to 348 and 315 OTUs in the USB and BGR treatments respectively (see Fig. 6a, b; ANOVA, *P* = 0.0007; t-test, *P* = 0.027, 0.002). Saprotroph OTU richness declined in all treatments relative to the control, though this effect was only significant in the BGR biochar treatment on balsam firs (ANOVA, *P* = 0.01). For balsam fir ECM fungal communities, both biochar treatments resulted in significant declines in Shannon diversity (ANOVA, *P* = 0.001; t-test, *P* = 0.006 and *P* = 0.08 for USB and BGR, respectively), which incorporates species evenness, while total fungal communities had similar Shannon diversity across all biochar treatments (see Fig. 6a, b). ECM and total fungal OTU richness on blue spruce roots were somewhat lower in biochar treatments than in the control, but differences were not significant. The decrease in ECM species richness and Shannon index on balsam fir indicates that biochar treatment favored ECM communities that were dominated by a low number of dominant species. ECM communities in all treatments were dominated by a single OTU identified as *Wilcoxina mikolae* which occurred in 49 out of 50 samples. *W. mikolae* increased in relative abundance (calculated as a percentage of ECM reads) from 60% in the control to 93% in the USB biochar treatment on balsam fir roots (ANCOM, *W*> 0.6). *W. mikolae* was similarly abundant in the USB biochar treatment on blue spruce at 92% of ECM reads, though it was also dominant in the blue spruce control treatment (83% of ECM reads) and thus the effect of USB biochar treatment was not significant. Two other OTUs from the Pyronemataceae (of which *Wilcoxina* is also a member) that were both classified as *Pulvinula* spp. were generally in low abundance (mean of 4%) but were dominant in three samples from the BGR treatment (36-84% relative abundance among ECM sequences), though they were observed too infrequently to test for differential abundance. Tests on the full fungal community indicated that the genera *Atractospora* and *Coprinellus* were in higher and lower abundance, respectively, in the fir BGR treatment compared to the control. Patterns in beta diversity show that total root-associated fungal communities were primarily structured by tree species (PERMANOVA, *P* = 0.0001), with little effect of biochar treatment (see Fig. 7a). Balsam fir and blue spruce total fungal communities were separated from each other along the first axis of the PCoA (see Fig. 7a). There was little effect of tree species on ECM community composition, but a significant effect of biochar treatment (see Fig. 7b; PERMANOVA, *P* = 0.0001), though this significant result may have been due to large differences in beta dispersion between groups (*P* < 0.0001).

**Figure 6.**
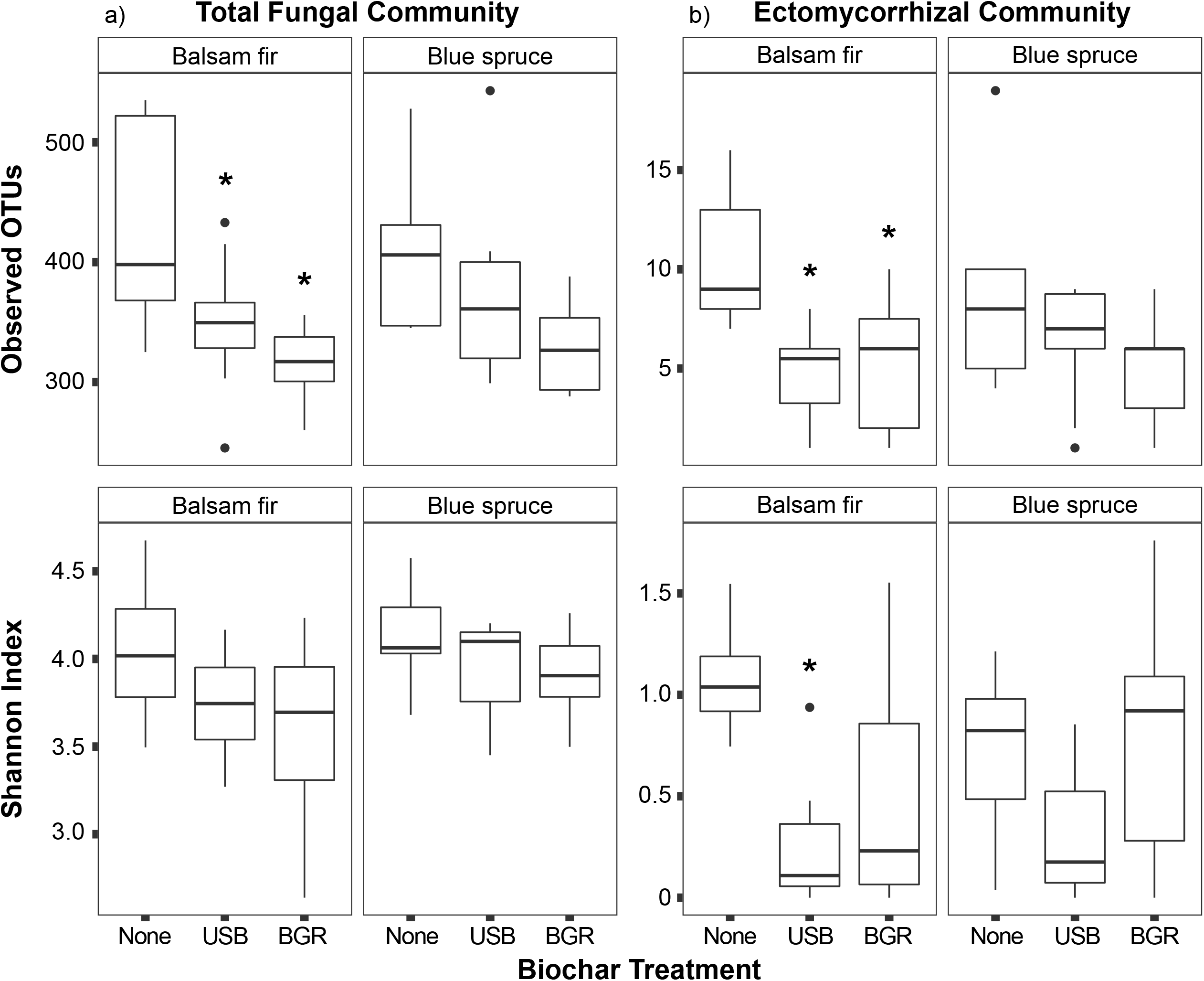
Patterns in root-associated fungal alpha diversity are plotted for the full fungal community (a) and the ectomycorrhizal fungal community (b). The number of observed OTUs and the Shannon Index were calculated as measures of alpha diversity. Error bars represent standard errors. Asterisks indicate significant differences between the biochar treatments and the control based on pairwise t-tests with Bonferroni-corrected p-values.

**Figure 7.**
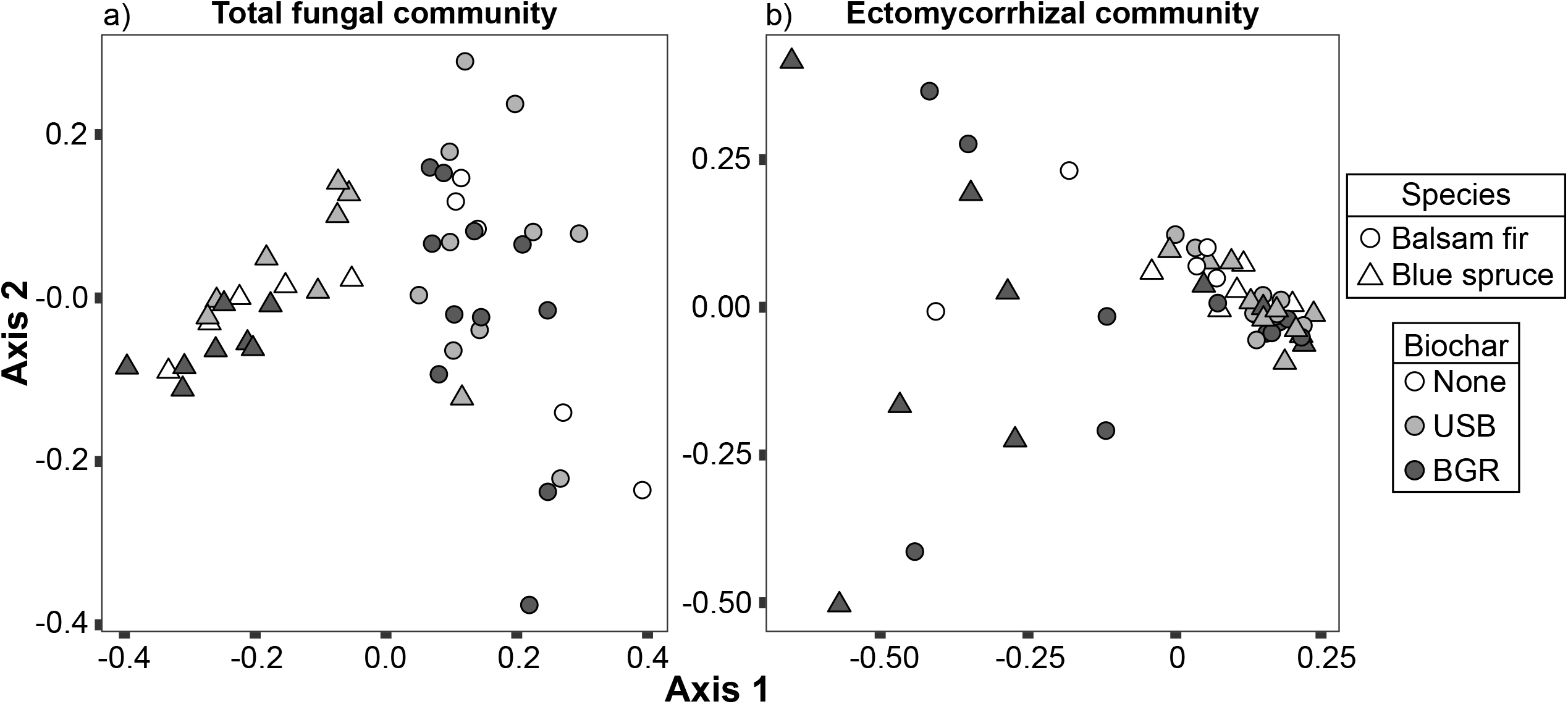
Principal coordinate analyses are plotted for the full fungal community (a) and the ectomycorrhizal fungal community (b) based on sequencing of ITS2 DNA amplicons prepared from root and rhizosphere samples. Points are colored by the biochar treatment and have different shapes based on the tree species.

## 4. Discussion

### 4.1 Mixed support of hypotheses

We hypothesized that biochar would 1) increase tree growth and survival, 2) improve soil physicochemical properties including soil moisture and nutrient concentrations, 3) improve soil microbial community functionality including extracellular enzyme activities (EEAs) and soil microbial biomass, and 4) stimulate ectomycorrhizal (ECM) colonization. As expected, our study found that biochar improved some physicochemical properties including soil moisture, phosphate availability, and the concentrations of multiple cations and also stimulated most EEAs. However, biochar also decreased nitrogen availability and ECM diversity, and reduced seedling growth and survival. Furthermore, there was no effect of biochar on soil microbial biomass as we had predicted. Other studies have found that biochar increases plant productivity by ameliorating unfavorable soil physicochemical conditions (Ali et al., 2017; Major et al., 2010). It is likely that the specific combination of biochars, crop species, and soil type were responsible for the negative outcomes that we observed. Thus, it is necessary to better understand the mechanisms of biochar’s effects on soil biology and plant productivity.

### 4.2 Differences between biochar types

Many of the soil variables in this study were more strongly affected by the BGR biochar than by the USB biochar, which raises questions about which properties of the biochars were responsible for their different effects. The BGR biochar had a lower N content than the USB biochar, which may have been responsible for a greater flux of DOC from the biochar. The lower N content in the BGR biochar is possibly related to its higher production temperature and pyrolysis time which could have resulted in greater N volatilization. Although biochar has often been considered to be very stable in soils, with turnover times on the order of decades to centuries, recent work has suggested that biochar actually contains a portion of C that is more labile and cycles on timeframes relevant to our three-year field experiment. Multiple studies have found that up to a few percent of the C present in biochar can be immediately extracted as DOC (Liu et al., 2019; Liu et al., 2015; Mukherjee and Zimmerman, 2013; Wu et al., 2018) and Hockaday et al. (2007) found that 0.20% of the total charred C in wildfire-derived charcoal was extractable as DOC 100 years after it had been produced, demonstrating that biochar leachates can contribute directly to the DOC pool in both the short- and long-term. Thus, it is possible that the biochars continued to act as a direct source of DOC throughout our three-year experiment. Alternatively, the positive effect of BGR biochar on DOC may have been due to its larger effect on pH than the USB biochar (see Fig. S3). A past study attributed positive effects of biochar on DOC to biochar’s effects on pH, suggesting that deprotonation of endogenous (not biochar-derived) DOC was responsible for the increase, rather than a direct contribution of biochar leachates to the DOC pool (Smebye et al., 2016). Despite this uncertainty, we can still conclude that BGR biochar had stronger effects on soil DOC pools than USB biochar, likely due to differences in biochar properties and interactions with soil.

### 4.3 Mechanisms of biochar’s effects on enzyme activities

The observed differences in DOC content between biochar treatments likely had cascading effects on soil microbial community functioning. Through structural equation modelling, we found that BGR biochar increased EEAs involved in N and P cycling (Fig. 2) by increasing DOC, though there was no DOC-mediated effect of biochar on EEA for any of the enzymes involved in C acquisition (see Fig. 3c). These findings suggest that biochar’s positive effects on EEAs due to increases in DOC may be more important for EEAs involved in nutrient acquisition than in C acquisition. The DOC mechanism may have contributed to the very large effects of biochar on LAP activity. Microbes can use biochar (Kuzyakov et al., 2009) and other aromatic C compounds (Boonchan et al., 2000; Hofrichter et al., 1999) as a C source. Thus, the increased DOC pool associated with BGR biochar may have induced microbial nutrient limitation by increasing the ratio of bioavailable C to N and P, which could have stimulated the production of enzymes that target those nutrients (Allison and Vitousek, 2005). This finding is supported by the decreases in both extractable (see Fig. S5) and resin-bound (see Fig. S4) inorganic N, which provides evidence for decreased N mineralization that may be due to DOC-induced N limitation. This mechanism was only observed in response to BGR biochar application because the USB biochar did not have an effect on DOC. However, there was only a single structural equation model (USB biochar’s effect on BG) in which biochar increased EEA by decreasing inorganic N. Thus, the positive effects of DOC on EEAs in BGR-treated soils could also be due to the DOC relieving C limitation in microbial enzyme synthesis. BGR biochar also increased the activities of all hydrolytic enzymes via its positive effect on soil moisture (see Fig. 3), which likely provided more favorable conditions for microbial metabolism and enzyme production (Xiao et al., 2018). A study by Wang et al. (2015) found that total N and exchangeable calcium are also potential controls on enzyme activities in biochar-amended soils. Other studies have found that biochar had no effects (Elzobair et al., 2016b) or negative effects (Wu et al., 2013) on extracellular enzyme activities.

Despite our findings that indirect effects mediated by DOC, moisture, and inorganic N are possible mechanisms for biochar’s effects on EEA, we still found that direct effects explained the greatest proportion of biochar’s effects on EEAs in all models (see Fig. 3 c, d). This suggests that the soil variables measured in our experiment did not fully capture the mechanisms of biochar’s effect on EEAs. The observed direct effects of biochar on EEAs could be explained by microbial community dynamics. Three hypotheses of biochar’s effects on EEA that involve the microbial community are possible: 1) increase of soil microbial biomass in response to biochar while enzyme production per unit microbial biomass remains constant, 2) alteration of soil microbial community composition, resulting in a higher EEA production per unit microbial biomass and 3) alteration of soil microbial community physiology, also resulting in a higher EEA production per unit microbial biomass. Hypothesis 1 can be ruled out because we did not observe changes in microbial biomass C in response to biochar application, in contrast to a past study that attributed biochar’s positive effect on extracellular enzyme activity to increased microbial biomass (Khadem and Raiesi, 2017). Hypotheses 2 and 3, however, represent viable explanation for biochar’s positive effects on EEAs. Hypothesis 2 is supported by significant alterations to root-associated fungal diversity and composition (see Figs. 6, 7). These changes were characterized by a decrease in total fungal, saprotroph, and ECM OTU alpha diversity on balsam fir trees in response to biochar (see Fig. 6). This pattern suggests that lower diversity fungal communities can be associated with higher rates of enzyme production. Lu et al. (2015) documented a similar pattern, finding that biochar decreased fungal diversity (measured by denaturing gradient gel electrophoresis), while increasing the activity of urease, invertase, and phosphatase. Hypothesis 3 is also certainly possible, though our data do not provide us with the necessary evidence to distinguish between microbial physiological and compositional mechanisms of biochar’s effects on EEAs.

### 4.4 Effects of biochar on fungal communities

We found that *Wilcoxina mikolae* sequences were present in almost all of our root samples, often as the dominant ECM species. USB biochar resulted in higher relative abundances of *W. mikolae* to the point of hyperdominance of ECM communities, which may have prevented colonization by other ECM fungi and hindered the development of a diverse ECM community. *Wilcoxina* spp. are considered to be pyrophilic (“fire loving”) ECM taxa because of their frequent occurrence on trees following wildfire (Jones et al., 2010). The frequent presence of *Wilcoxina* in recently burned stands has been attributed to both its affinity for disturbance and the thermotolerance of its propagules (Baar et al., 1999). However, our data provide evidence that in addition to previously demonstrated linkages with disturbance and soil heating, the presence of black C in soil may be another factor that favors *Wilcoxina*. This hypothesis is supported by the study of Robertson et al. (2012) who found that an E-strain fungus (likely *Wilcoxina*) increased in colonization in response to biochar and constituted that majority of ECM root tips when 10% biochar was amended into soil. Our finding that *Wilcoxina* sequence abundance is increased by USB biochar suggests that effects of biochar application on microbial community composition may be relevant to understanding the assembly of microbial taxa following wildfires or prescribed fires in non-agricultural systems, thus providing a linkage between the role of black C role in natural and agricultural systems. *Wilcoxina* is classified as an ectendomycorrhizal fungus, meaning that it colonizes intracellularly in addition to the characteristic extracellular Hartig net and mantel. The positive responses of *Wilcoxina* to biochar may be due to biochar’s liming effect as ectendomycorrhizal fungi are known to form abundant mycorrhizae at slightly alkaline soil pH (Danielson and Pruden, 1989; Taylor and Finlay, 2003; Theodorou and Bowen, 1969), while most ECM fungi prefer moderately acidic soils (Erland et al., 1990; Hung and Trappe, 1983).

The negative effects of biochar on ECM diversity in balsam fir (see Fig. 6b) may explain the greatly decreased tree growth, which was not observed in blue spruce (see Fig. 5c, d). Although results are mixed from studies measuring the effect of ECM diversity on tree growth and nutrient uptake, there is promising evidence that ECM diversity has positive effects on plant fitness (Baxter and Dighton, 2001; Jonsson et al., 2001; Köhler et al., 2018). Additionally, the high abundance of *Wilcoxina* sequences may have hindered tree performance as Mikola (1988) found that in low nutrient conditions, *W. mikolae* decreased Scots pine growth relative to both uninoculated control seedlings and seedlings inoculated with forest humus which presumably contained diverse ECM communities.

### 4.5 Effects of biochar on tree performance and weed productivity

The reduced survival and growth of seedlings in response to BGR biochar may have been due to the negative effects of the biochar on N availability, as we observed decreases in inorganic N and increases in the DOC:TDN ratio relative to the control. Although conifer seedlings responded negatively to BGR biochar, this is not evidence of a general inhibitory effect of biochar on plant growth, because we observed positive effects of BGR biochar on weeds, with up to a 94% increase in weed biomass. This contrast between the responses of the weeds and the tree seedlings to biochar is similar to the findings of a meta-analysis of the effect of biochar on plant productivity (Biederman and Harpole, 2013), which found that on average annual plants responded positively to biochar, while perennial plants responded slightly negatively (though non-significantly) to biochar application. This increase in weed growth in the biochar plots may have contributed to the decrease in ECM fungal diversity as fast-growing herbaceous plant species have been previously shown to decrease ECM colonization of conifers (Wolfe et al., 2008). Higher weed biomass likely also exerted a competitive effect on the tree seedlings for water, nutrients, and light and may be related to the decrease in inorganic N. The problem of competition from weeds may have been compounded by interference of biochar with the pre-emergent herbicides that we applied. Multiple experiments have shown that biochar can interfere with the efficacy of pre-emergent herbicides (Graber et al., 2012; Spokas et al., 2009; Yu et al., 2006; Zheng et al., 2010) which could increase competition for water and nutrients from herbaceous weeds. Along with another study that found dramatic increases in weed cover in biochar-amended plots (Major et al., 2005), our study demonstrates that biochar can have the unintended consequence of increasing pressure from weeds. It is also worth noting that herbicide application may have affected microbial processes including EEAs and mycorrhizal colonization. However, our goal was to manage this experiment similarly to a typical Christmas tree plantation and because herbicides are widely used in the industry (Landgren et al. 2003), they should be considered as part of the agroecosystem.

Although our study found that biochar negatively affected the performance of conifers in a Christmas tree plantation on well-drained Marlette fine sandy loam soil, we observed wide ranging effects on soil physical, chemical, and biological properties that could be harnessed to improve conditions in other systems with different combinations of crops and soil types. When applied to our soils, which were already close to neutral in pH, biochar resulted in a soil pH that was higher than the optimal level for the growth of blue spruce and balsam fir (Hart et al., 2009), but this increase in pH may actually be beneficial in systems that have more acidic soil, which is often the case for Christmas tree plantations that have been through multiple rotations and have been subject to the acidifying effects of conifer litter and fertilizer (Hornung, 1985). Indeed, a recent meta-analysis found that soils with a lower initial pH were more likely to result in greater crop yields with biochar application (Jeffery et al., 2017). Both biochars also exhibited the potential to sequester heavy metals (see Fig. S4), which could improve tree performance in contaminated soils or reduce tree performance if availability is reduced to levels less than optimal for plant growth. Likewise, the increase we observed in soil moisture with biochar application may have greater importance in more arid climates where soil moisture is a stronger constraint on tree growth and survival. Further investigations may provide a predictive framework for understanding which types of biochar will result in soil physicochemical closest to the optima of a given crop species or desired microbially mediated process (e.g. nitrification, mycorrhizal colonization), opening up the possibility for “designer biochars” to ameliorate specific soil quality issues (Novak et al., 2014).

## 5 Conclusions

Despite the overwhelming interest in biochar application as a strategy to increase crop productivity and modify soil biological processes, there are still relatively few studies on woody crops, and even fewer on those that associate with ectomycorrhizal fungi. Our study, which is one of the most comprehensive investigations into biochar’s effects on an ectomycorrhizal cropping system, provides strong evidence that a much greater understanding of the effects of biochar on soil biology and crop performance is needed before biochar is applied widely in cropping or forestry systems with ectomycorrhizal plants. Our findings of increased extracellular enzyme activity, lower ectomycorrhizal fungal diversity, decreased tree performance, and modifications to a number of soil physicochemical properties suggests that complex interactions between plants, microbes, and soils need to be considered when predicting the effects that biochar will have on a given cropping system. Despite these possible pitfalls, the potential for biochar to be economically viable in woody cropping is large because of the relatively long rotation periods which may justify the greater upfront investment in biochar if there are proven benefits. A greater mechanistic understanding of the context-dependencies that determine biochar’s effects on crop performance will help develop science-based recommendations on biochar application for growers.

## Acknowledgments

We thank Randy Klevickas and Paul Bloese for management of the field experiment. We thank Chase O’Neil, Chase Kasmerchak, Eion Riley, Kristen Przano, Ethan Rocklin, and Charity Moore for assistance in field sampling and laboratory work. We thank Reid Longley for preparing DNA libraries and doing initial sequence data processing. We thank Darian Smercina and Andrew Curtright for advising on certain procedures. We thank Biogenic Reagents for providing BGR biochar. This project was supported by the Michigan Christmas Tree Association, Michigan State University Project GREEEN, the Michigan Department of Agriculture and Rural Development Specialty Crop Block Grant Program, and by the USDA National Institute of Food and Agriculture, McIntire Stennis project 1006839.

## Figure captions

**Table 1.** Concentrations of elements in BGR and USB biochars are reported with all values in percent mass.

**Supplemental file 1.** Contains figures S1-S5 and tables S1, S2

**Supplemental file 2.** Contains a FASTA file of representative sequences of OTUs generated by amplicon sequencing of the fungal ITS2 gene region. FASTA headers match OTU IDs in supplemental files 3, 4, and 5.

**Supplemental file 3.** Contains taxonomic assignments of OTUs generated by the feature-classifier from QIIME trained on the UNITE database.

**Supplemental file 4.** Contains the full OTU table of the fungal ITS2 region, with OTU counts rarefied to an even sequence depth of 13472. OTU IDs match those in supplemental files 2 and 3 and sample names match those in supplemental file 6.

**Supplemental file 5.** Contains the data matrix of OTUs identified as ectomycorrhizal by FUNGuild with OTU counts relativized. OTU IDs match those in supplemental files 2 and 3 and sample names match those in supplemental file 6.

**Supplemental file 6.** Contains the sample metadata for root samples that had fungal ITS2 amplicons sequenced. Sample names match those in supplemental file 3.

## Abbreviations

EEA: extracellular enzyme activity
ECM: ectomycorrhizal
TDN: total dissolved nitrogen
DOC: dissolved organic carbon
PRS: plant root simulator
BG: β-glucosidase
CBH: cellobiohydrolase
LAP: leucine aminopeptidase
NAG: β-N-acetyl glucosaminidase
PER: peroxidase
PHEN: phenoloxidase
PHOS: acid phosphatase
EDX: energy dispersive X-ray spectroscopy
ITS: internal transcribed spacer
PCR: polymerase chain reaction
ANOVA: analysis of variance
PERMANOVA: permutational multiple analysis of variance
ANCOM: analysis of composition of microbiomes
OTU: operational taxonomic unit

