## Supplemental File 1 for "Biochar alters soil properties, microbial community diversity, and enzyme activities, while decreasing conifer performance"

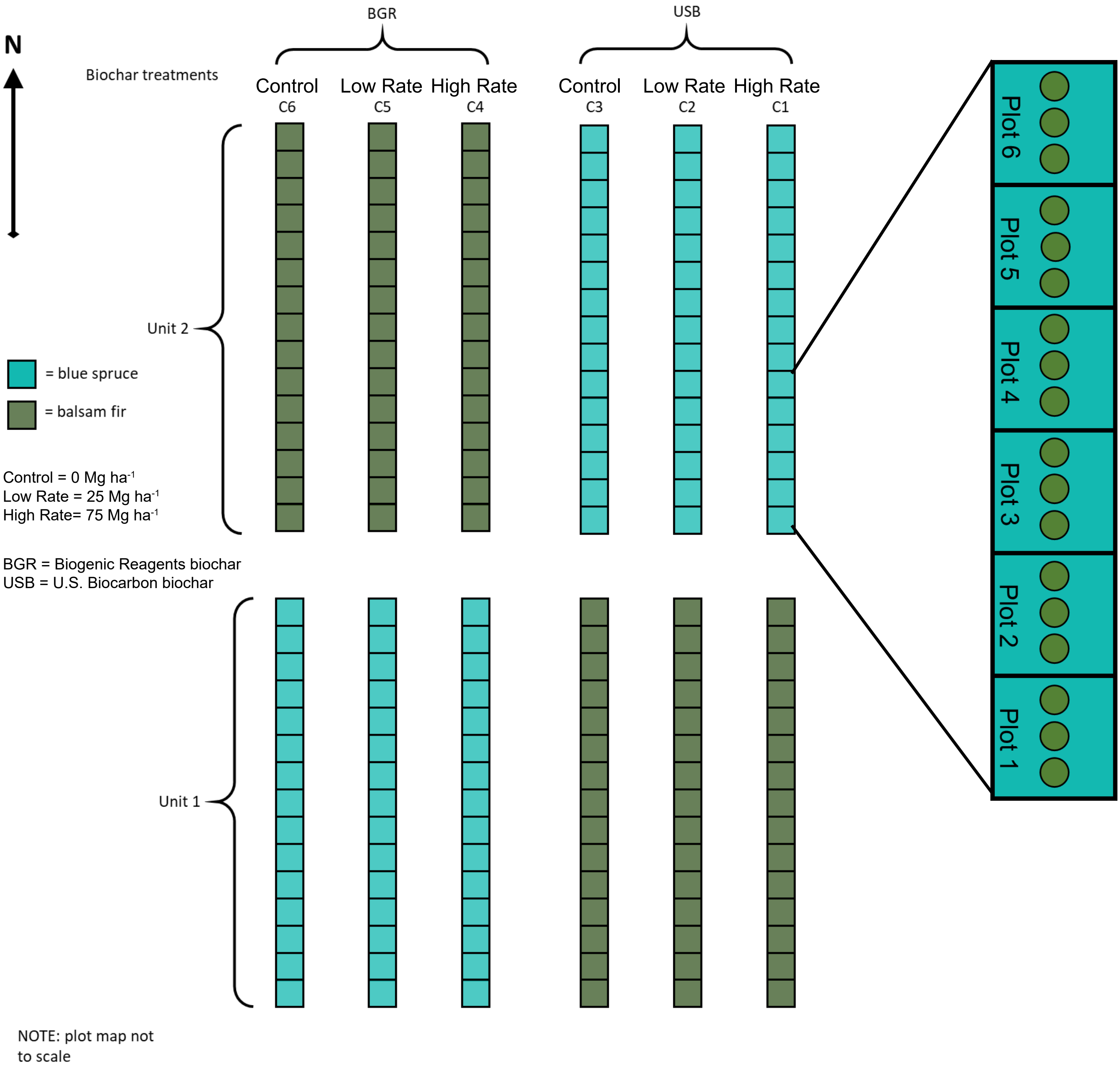

Supplemental figure S1. The layout of the field experiment. The experimental planting was divided into two units, each of which contained six columns. Each of the columns represented a single treatment unit. There were 15 plots within each column that all received the same treatment. Each plot was planted with three tree seedlings (represented by green circles). BGR = Biogenic Reagents biochar. USB = U.S. Biocarbon biochar.

| **2016** | *P* |
|---|---|
| Treatment | <0.0001 |
| Date | <0.0001 |
| Treatment:Date | 0.017 |
| Marginal R² | 0.77 |
| Conditional R² | 0.77 |

| **2017** | *P* |
|---|---|
| Treatment | <0.0001 |
| Date | <0.0001 |
| Treatment:Date | 0.503 |
| Marginal R² | 0.69 |
| Conditional R² | 0.72 |

| **2018** | *P* |
|---|---|
| Treatment | <0.0001 |
| Date | <0.0001 |
| Treatment:Date | 0.601 |
| Marginal R² | 0.75 |
| Conditional R² | 0.76 |

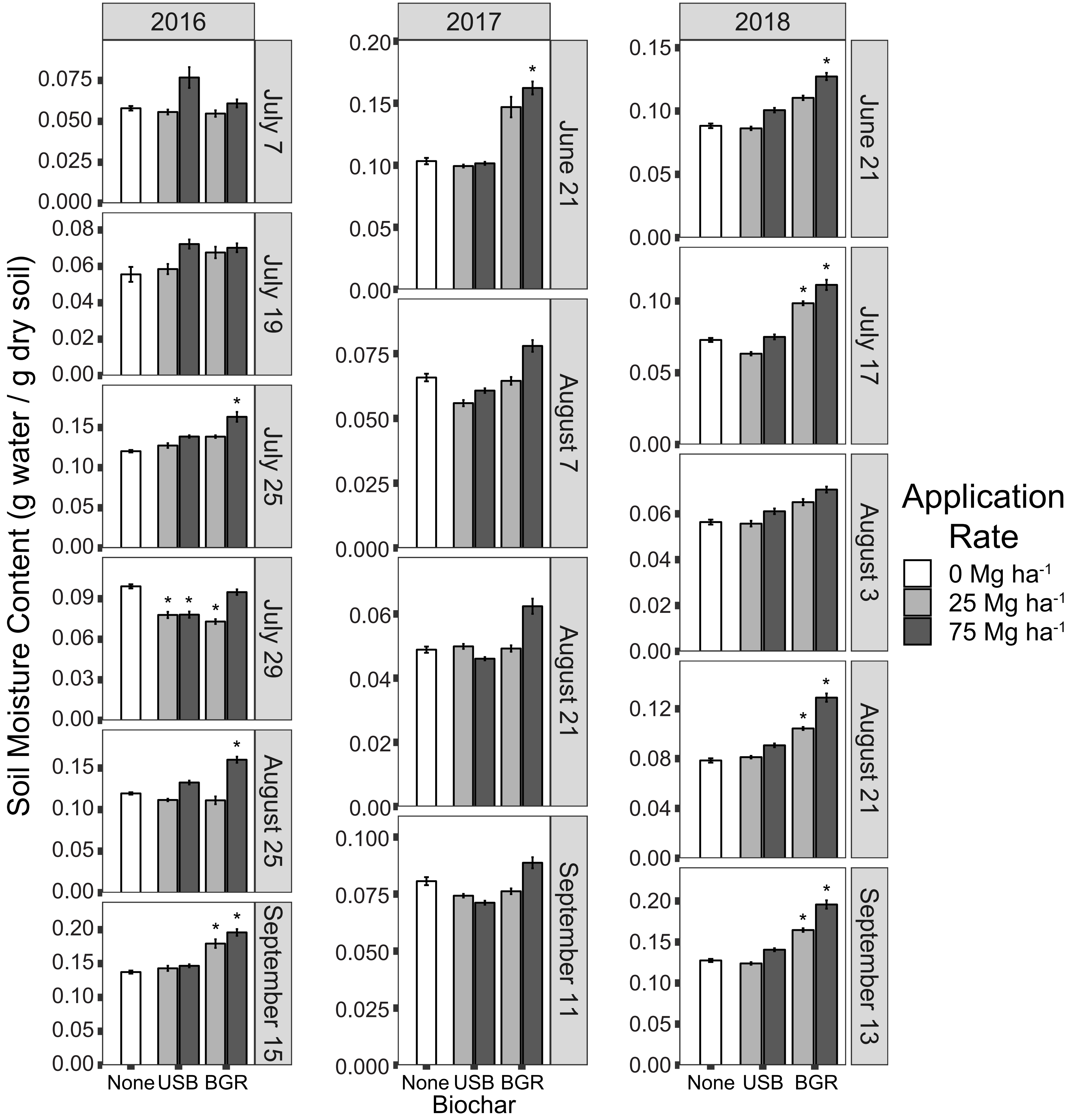

Supplemental figure S2. This figure presents soil moisture data collected across 2016, 2017, and 2018. Separate mixed models were fit to each year, which are summarized at the top of each column. Asterisks in the panels indicate significant ($P < 0.05$) differences between biochar treatments and the control for each sampling date after Bonferroni corrections assuming four comparisons. Error bars represent standard errors

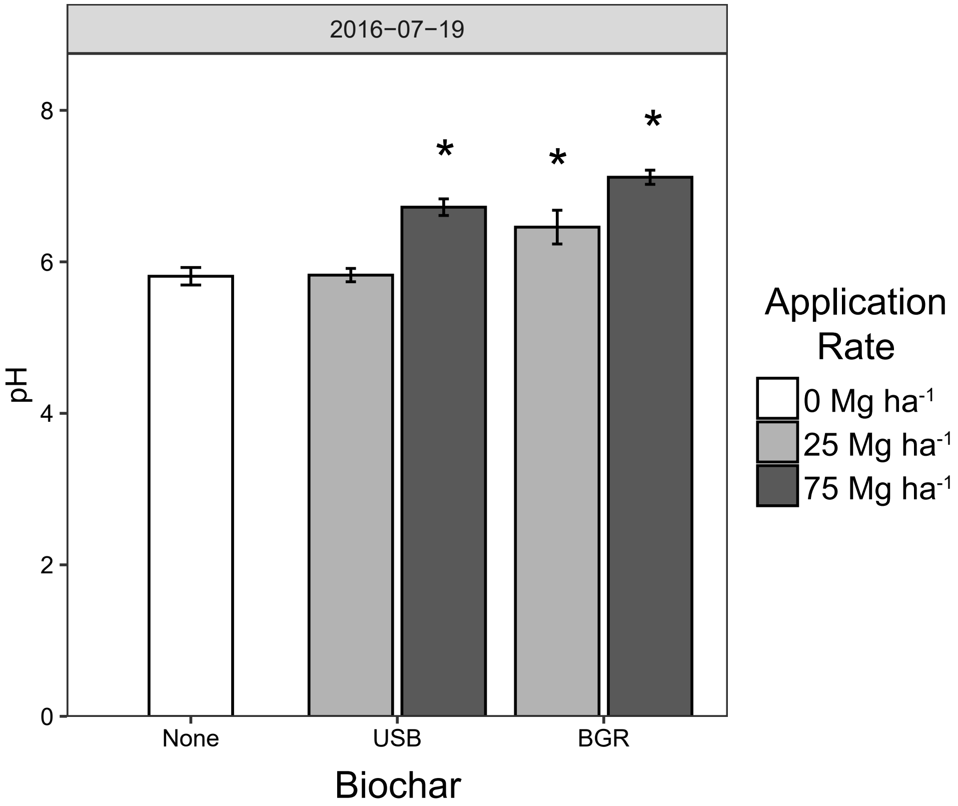

Supplemental figure S3. Mean pH values of soil (measured in a 2:1 suspension with distilled water) from each treatment are plotted from a sampling date in the first growing season. Error bars represent standard errors. Asterisks represent significant differences between the biochar treatments and the control treatment based on pairwise t-tests with Bonferroni correction of p-values ($P < 0.05$).

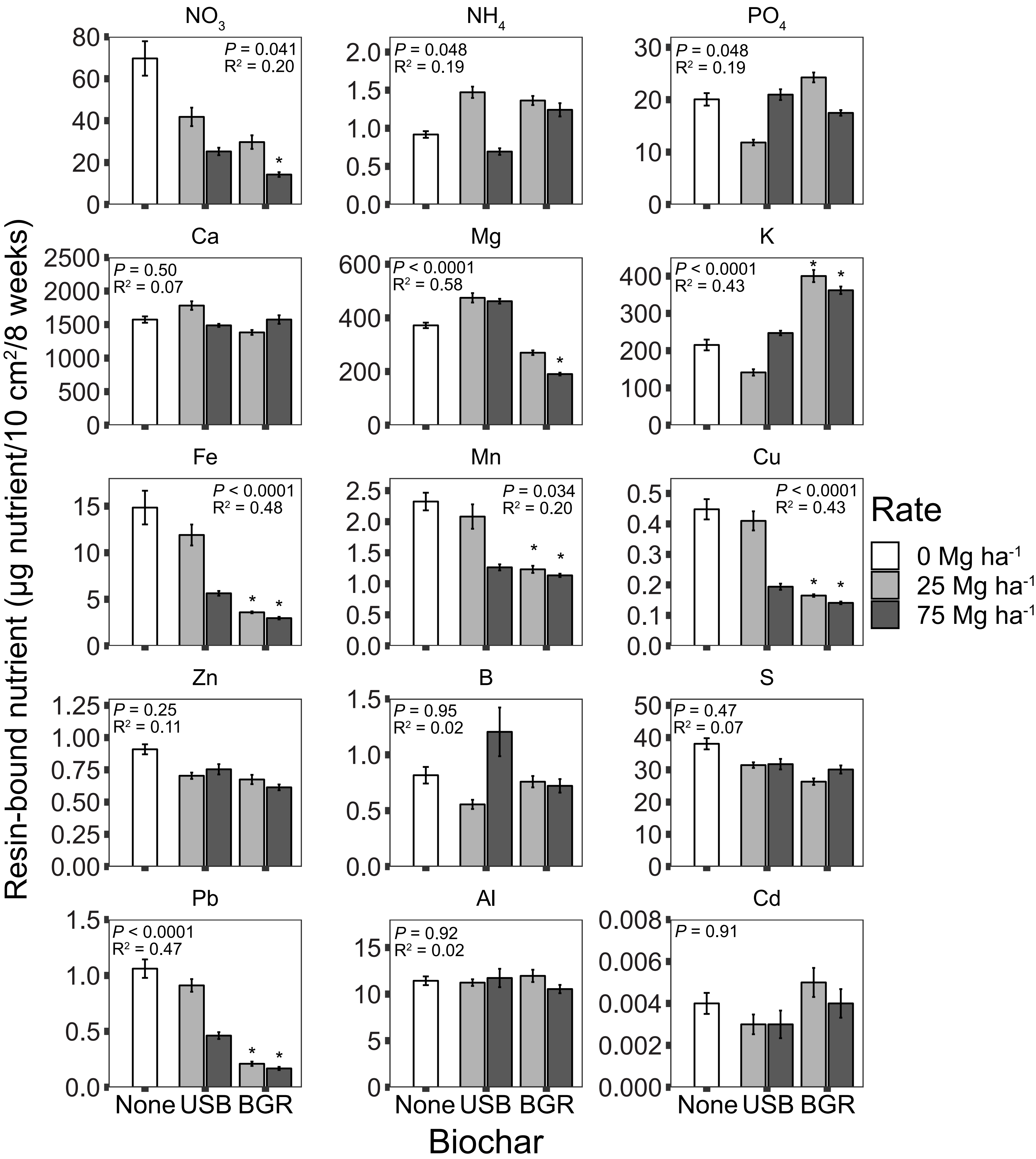

Supplemental figure S4. The bioavailabilities of 15 ions were monitored over 8 weeks in the 2018 growing through field-installation of Plant Root Simulator resin probes.Test statistics from ANOVAs tests are displayed in each panel, while asterisks indicate significant pairwise differences between biochar treatments and the control ($P$ < 0.05) based on t-tests with Bonferroni corrected $P$-Values. Error bars represent standard errors

| 2016 | *P* |
|---|---|
| Treatment | <0.0001 |
| Date | <0.0001 |
| Treatment:Date | <0.0001 |
| Marginal $R^2$ | 0.80 |
| Conditional $R^2$ | 0.80 |

| 2017 | *P* |
|---|---|
| Treatment | <0.0001 |
| Date | <0.0001 |
| Treatment:Date | <0.0001 |
| Marginal $R^2$ | 0.97 |
| Conditional $R^2$ | 0.97 |

| 2018 | *P* |
|---|---|
| Treatment | 0.003 |
| Date | <0.0001 |
| Treatment:Date | <0.0001 |
| Marginal $R^2$ | 0.52 |
| Conditional $R^2$ | 0.58 |

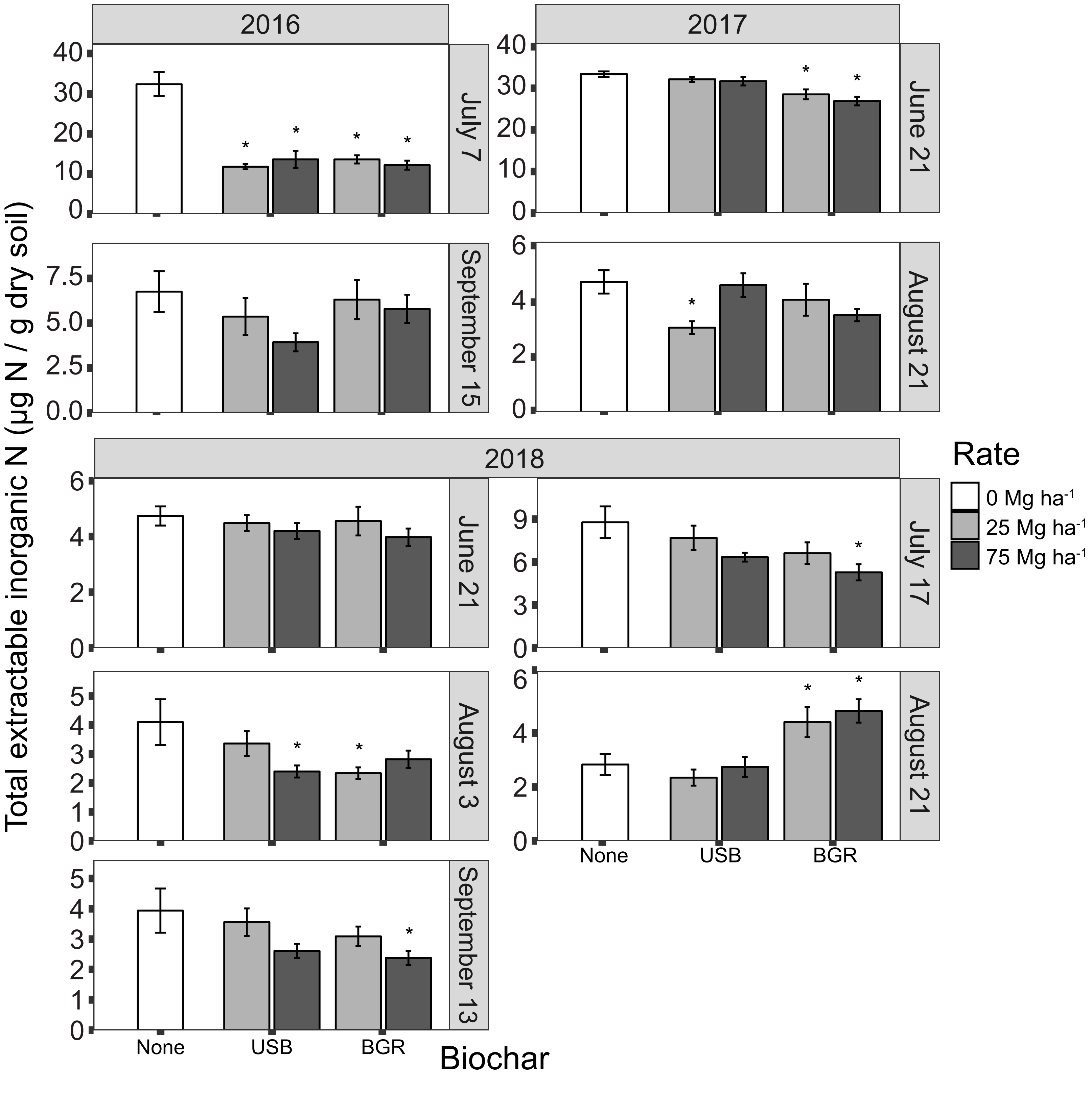

Supplemental figure S5. Total extractable inorganic N (sum of $NH_4$ and $NO_3$) is plotted for sampling dates across 2016, 2017, and 2018 with error bars depicting standard errors. Separate mixed models were fit to each year, which are summarized at the top of each column. Asterisks in the panels indicate significant ($P < 0.05$) differences between biochar treatments and the control for each sampling date based on t-tests with Bonferroni corrected p-values. Note the variation in y-axis scale, particularly for July 7, 2016 and June 21, 2017.

| Date | Time | Product Name | Active Ingredient | App Type | Application Rate | Comments |
|---|---|---|---|---|---|---|
| 13-May-16 | 10:00 AM | Simazine 90DF | Simazine | 3' band | 3.36 kg/ha | w/ SPSO backpack and 8006 nozzle |
| 13-May-16 | 10:00 AM | Pendulum Aquacap | Pendimethalin | 3' band | 3.4 L/ha | w/ SPSO backpack and 8006 nozzle |
| 29-Jun-16 | 10:00 AM | Kill-Zall II | Glyphosate | directed band | 4.2 L/ha | Applied 2' band on either side of trees w/ OC06 nozzle and SPSO backpack sprayer |
| 19-Sep-16 | 1:30 PM | Weedar 64 | 2,4-D | 3' band | 0.83 L/ha | Hand sprayed w/ SPSO backback sprayer & 8006 nozzle. |
| 19-Sep-16 | 1:30 PM | Kill-Zall II | Glyphosate | 3' band | 0.63 L/ha | Hand sprayed w/ SPSO backpack sprayer & 8006 nozzle. |
| 12-May-17 | 2:00 PM | Envoy | Clethodim | 3' band | 0.46 L/ha | SPSO backpack w/ 8006 flat fan nozzle. Total of 3.5 gal @ 0.5 oz/gal. |
| 26-Apr-18 | 11:30 AM | Simazine 90DF | Simazine | 3' band | 96 g/ha | SPSO sprayer, 8006E nozzle; 60 F, NW wind at 5 mph. 3.3 g/gal = 3 lb/a. |
| 26-Apr-18 | 11:30 AM | Sureguard | Flumioxazin | 3' band | 4.3 mL/ha | SPSO sprayer, 8006E nozzle, 60F, NW wind at 5 mph. 1 tsp/gal = 10 oz/a. |
| 26-Apr-18 | 11:30 AM | Kill-Zall III | Glyphosate | 3' band | 0.68 L/ha | SPSO sprayer, 8006E nozzle, 60 F, NW wind at 5 mph. 1qt/a = 0.8oz / gal |

Table S1. The record of herbicide applications throughout the experiment.

| | Analysis Performed | | | | | | | | | | |
|---|---|---|---|---|---|---|---|---|---|---|---|
| Date | U of Maine Testing Lab | Extracellular enzyme Activity | Inorganic nitrogen | Dissolved organic matter | Bulk density | Scanning electron microscopy | Soil moisture | Tree growth and mortality | pH | Plant root simulators | Root amplicon sequencing |
| 6/23/2016 | | | | | | | | x | | | |
| 7/7/2016 | | x | x | | | | x | | | | |
| 7/19/2016 | | x | | | | | x | | x | | |
| 7/25/2016 | | x | | | | | x | | | | |
| 7/29/2016 | | x | | | | | x | | | | |
| 8/25/2016 | x | x | | | | | x | | | | |
| 9/15/2016 | | x | x | | | | x | | | | |
| 9/23/2016 | | | | | | | | x | | | |
| 6/21/2017 | | x | x | x | | | x | x | | | |
| 7/17/2017 | | x | | | | | | | | | |
| 8/7/2017 | | x | | | | | x | | | | |
| 8/21/2017 | | | x | | | | x | | | | |
| 9/11/2017 | | x | | | | | x | | | | |
| 9/17/2017 | | | | | | | | x | | | |
| 10/26/2017 | | | | | | | | | | | x |
| 6/21/2018 | | x | x | x | | | x | | | | |
| 7/10/2018 | | | | | | | | x | | | |
| 7/17/18 to 9/13/18 | | | | | | | | | | x | |
| 7/17/2018 | | x | x | x | | | x | | | | |
| 8/3/2018 | | x | x | x | | | x | | | | |
| 8/21/2018 | | x | x | x | | | x | | | | |
| 9/13/2018 | | x | x | x | | | x | | | | |
| 10/1/2018 | | | | | x | x | | | | | |
| 10/8/2018 | | | | | | | | x | | | |

Table S2. A record of sampling dates for soil, plant growth, and microbial data collection. Note that the first column labelled "U of Maine Testing Lab" refers to soil tests done at the University of Maine for the following properties: soil pH, loss on ignition, effective cation exchange capacity, Ca, K, Mg, P, Al, B, Cu, Fe, Mn, Na, S, and Zn.
